# Seasonal variation in sexual readiness in a facultatively sexual freshwater cnidarian with diapausing eggs

**DOI:** 10.1101/2020.05.27.119123

**Authors:** Jácint Tökölyi, Réka Gergely, Máté Miklós

## Abstract

Facultative sexuality combines clonal propagation with sexual reproduction within a single life cycle. Clonal propagation enables quick population growth and the occupancy of favorable habitats. Sex, on the other hand, results in the production of offspring that are more likely to survive adverse conditions (such as the resting eggs of many freshwater invertebrates). In seasonal environments, the timing of sex is often triggered by environmental cues signaling the onset of winter (e.g. temperature drop or changes in photoperiod). Organisms switching to sex to produce resting eggs under these conditions face a trade-off: responding too early to an environmental cue increases the chances of missing out in clonal propagation, while having a delayed response to deteriorating conditions entails the risk of parental mortality before sexual reproduction could be completed. To mitigate these risks, increased sensitivity towards environmental cues with the onset of the winter might be an adaptive strategy. To test this hypothesis, we investigated sexual propensity and time to gonadogenesis in clonal strains derived from spring- and autumn-collected polyps of *Hydra oligactis*, a facultatively sexual freshwater cnidarian where sex only occurs prior to the onset of winter. We show that autumn-collected individuals and their asexual offspring have a higher propensity for sex and require less time for gonad development compared to strains established from spring-collected individuals that were kept under similar conditions in the laboratory. To see if the above results can be explained by phenotypic plasticity in sexual readiness, we exposed cold-adapted lab strains to different lengths of warm periods. We found that sexual propensity increases with warm exposure. Our results suggest that reciprocal cold and warm periods are required for sex induction in *H. oligactis*, which would ensure proper timing of sex in this species. Increased sensitivity to environmental deterioration might help maximize fitness in environments that have both a predictable (seasonal) and an unpredictable component.

## Introduction

Facultatively sexual organisms combine clonal reproduction with occasional sex and show astonishingly diverse life histories (Stelzer and Lehtonen 2016, Kokko 2020). The switch from asexual to sexual reproduction in these organisms can be highly variable within and between species (e.g. Tessier and Cáceres 2004, Navarro et al. 2013, Franch-Gras et al. 2017, Ryan and Miller 2019). Central to research on facultatively sexual organisms is to identify factors that drive this variation.

A common feature in the ecology of facultatively sexual animals is that many of them inhabit ephemeral or highly seasonal environments where favorable periods alternate regularly or irregularly with unfavorable periods. Favorable periods are often associated with clonal growth which enables exploitation of available habitat patches and the resource boom (Hadany and Otto 2009, Stelzer 2012, Stelzer and Lehtonen 2016). Unfavorable periods, on the other hand, tend to elicit sexual reproduction and result in diapausing stages, such as the resting eggs of aphids, monogonont rotifers, water fleas and hydras (Simon et al. 2002, Tessier and Cáceres 2004, Schröder 2005, Steele et al. 2019). Because of the strong association between sex and diapause in facultatively sexual species, variation in sexual investment needs to be considered in the context of environmental fluctuations and selection for diapause (Tessier and Cáceres 2004, Stelzer and Lehtonen 2016).

In the seasonal environment of the temperate zone environmental conditions change predictably each year and result in the alternation of favorable and unfavorable periods. Unfavorable periods can occur either during the summer (e.g. if drought is a major source of mortality) or in the winter (if freezing occurs). In either case, the regular changes in the environment characteristic of seasonal habitats provides cues to detect the deterioration of the environment and to trigger the switch to a diapausing stage. For instance, in case of water fleas changes in food, photoperiod and population density induce the parthenogenetic production of males, which elicit sexual mode of reproduction and the production of resting eggs in females (Tessier and Cáceres 2004, Camp et al. 2019). Decreasing day length and temperature induce a switch from parthenogenetic to sexual reproduction in aphids (Simon et al. 2002), while in hydra both increases and decreases in temperature can be signals that initiate gametogenesis, depending on the species (Reisa 1973).

While seasonally varying environmental cues can provide reliable information about environmental change, no environment is entirely predictable. Therefore, the correlation between environmental cues and environmental change is not perfect. In such a situation, organisms face a trade-off between responding early and losing the opportunity for asexual reproduction if the environment does not change as expected or responding late and thereby risking increased mortality if the environment does change. To mitigate these risks, increased sensitivity toward sex-inducing environmental stimuli with the progress of seasons might be an optimal strategy in an environment that is both seasonal and unpredictable. Indeed, in monogonont rotifers the propensity to produce sexual females in response to an environmental cue (crowding) is low after hatching from a diapausing egg but increases following several generations of asexual reproduction (Schröder and Gilbert 2004). In case of *Daphnia* species, information about conditions experienced by parthenogenetic mothers (food and daylength) can be transmitted to the offspring, which ensures correct timing of resting egg production in the latter (Alekseev and Lampert 2001). Likewise, in hydra strains kept in the laboratory under stable conditions, propensity for sex is low after establishing strains from a polyp that hatched from an egg, but increases during several years of asexual culture (Noda 1982). These examples suggest that sex induction in response to environmental cues in facultatively sexual species can be much more complex than a simple stimulus-response. However, relatively little is known about how sensitivity to environmental cues changes during the life cycle of facultatively clonal species and the mechanisms behind this phenomenon.

The freshwater cnidarian *Hydra oligactis* inhabits highly seasonal environments of the temperate zone. *H. oligactis* polyps reproduce asexually throughout much of the year but switch to sexual reproduction in response to cooling (Reisa 1973). Throughout the distribution range sexual reproduction in natural habitats occurs from late summer to December (Welch and Loomis 1924, Ribi et al. 1985, Sebestyén et al. 2018). Sexual reproduction results in the production of diapausing eggs that tolerate desiccation and freezing (Steele et al. 2019). Sexual reproduction appears to be highly costly to the adults. Polyps that produce gametes have reduced numbers of interstitial stem cells and nematocytes necessary for food capture, they have substantially impaired regeneration capacity, lose their ability to feed and ultimately experience high mortality (Yoshida et al. 2006, Sebestyén et al. 2018, Tomczyk et al. 2020).

Given these apparent costs of sexual reproduction in *H. oligactis*, it is perhaps not surprising that not all polyps reproduce sexually even if exposed to the same environmental stimulus. In the wild, only a subset of the population reproduces sexually at any time during the autumn (Welch and Loomis 1924, Ribi et al. 1985, Sebestyén et al. 2018). Strains derived from the same population and kept under standard conditions in the laboratory differ in their propensity for sex (Tökölyi et al. 2017b). Even within a strain derived through asexual propagation of a single polyp, there is variation in sex induction capacity in response to the same environmental cue (cooling; Sebestyén et al. 2019).

Here, we asked if sexual readiness of *H. oligactis* exposed to the same environmental conditions shows seasonal variation. Sexual reproduction and production of resting eggs in this species only occurs prior to the onset of winter (Reisa 1973). Therefore, maintaining high preparedness throughout the year might not be optimal, especially if preparedness is costly (Tökölyi et al. 2012). Hence, *H. oligactis* polyps might be expected to show increased levels of preparation as the onset of winter approaches.

To test our hypothesis, we collected hydra strains from a single population during spring and autumn in two years (four collections in total) and established strains from them. These strains were kept under standard conditions in the laboratory and were induced to produce gametes by simulating the onset of winter via lowering the temperature in the laboratory. We recorded the presence of sexual reproduction as well as time to gonadogenesis to estimate sexual readiness of polyps and compared these variables across seasons. Furthermore, to see whether differences in sexual readiness between seasons is due to phenotypic plasticity, as predicted by our hypothesis, we performed warm exposure experiments in two lab strains (one male and one female) and looked at changes in sexual readiness in response to exposure to elevated temperature.

## Methods

### Field collection

Strains were established from a single lake in Central Hungary (Tiszadorogma, 47.67 N, 20.87 E) on four dates: 31^st^ May 2018 (henceforth called Spring 2018), 1^st^ October 2018 (Autumn 2018), 16^th^ May 2019 (Spring 2019) and 24^th^ September 2019 (Autumn 2019). The collection site is a small, shallow oxbow lake directly connected to the Tisza river with a small canal. The *H. oligactis* population inhabiting this lake was the subject of previous studies aimed at explaining life history variation in *Hydra* (Tökölyi et al. 2017a, 2017b, Sebestyén et al. 2018, Miklós et al. 2019).

On each collection occasion hydra polyps were collected from submerged vegetation along the shoreline from several distinct locations that were at least 2 m from each other. Polyps found on each location were put in a Falcon tube in lake water and brought to the laboratory in a cool box on the day of collection.

### Lab maintenance of strains and induction of sex

Hydra polyps brought to the laboratory were immediately moved to hydra medium (M-solution: 1mM Tris, 1mM NaCl, 1mM CaCl2, 0.1mM KCl, 0.1mM MgSO4 at pH 7.6; Lenhoff 1983). We selected up to five polyps from each location to establish strains from them through asexual propagation. Both wild-collected polyps and their asexual offspring were kept individually in 6-well plates, with 5 ml M-solution per well. They were fed twice per week with 20 μl suspension of freshly hatched Artemia nauplii (see for a description of the feeding method Tökölyi et al. 2016) and moved to fresh hydra medium approximately 1 hour after each feeding.

The asexual propagation phase lasted for 10 weeks. During this time polyps were kept in a climate chamber at 18 °C and a 12/12 h light-dark cycle. Asexual polyps produced during this period were retained and moved to empty wells with fresh medium in the plates. To keep samples at a manageable size, only two buds/week were retained in strains with a high budding rate and the maximum number of polyps/strain was set to N=18 (i.e. three 6-well plates).

After 10 weeks both wild derived polyps and their lab-derived asexual offspring were moved to a thermostatic cooler with 8 °C and a 8/16 h light-dark cycle, to simulate the onset of winter and induce gametogenesis (henceforth called “cold phase”). Asexual offspring produced after this time were no longer retained for collecting phenotypic data. Experimental animals were kept for five months under these conditions. During the cold phase polyps were checked twice per week under a stereo microscope (Euromex, Stereoblue) to detect the start of gonadogenesis.

### Warm exposure experiment

Two strains of *H. oligactis* were used to investigate the effect of warm exposure on sexual propensity: one male (C2/7) and one female (X11/14) strain. Both originate from the same population in Tiszadorogma but were collected earlier (autumn 2016) and kept in the lab on 18 °C and 12/12 h light/dark cycle since then (see Sebestyén et al. 2019 for a description of these two strains).

To test the effect of seasonal changes in temperature and warm exposure on sexual readiness in *H. oligactis*, we first generated “cold strains” from C2/7 and X11/14. Cold strains are hydra polyps asexually propagated on 8 °C. While lowering the temperature from 18 °C to 8 °C induces sex in most adults, they often produce a few buds before gonadogenesis inhibits budding. These asexual offspring have reduced propensity for sex and a higher asexual rate even if kept on 8 °C with the adult polyp (Sebestyén et al. 2019). By asexually propagating these buds, a large number of asexual offspring can be generated on 8 °C (cold strains). We established cold strains by selecting adult polyps from C2/7 and X11/14 kept on 18 °C (N=19 from both), moving them to 8 °C and asexually propagating buds that detached from their sexual parents on 8 °C.

After a large number of asexual offspring were obtained, cold strains were divided into three groups: (1) control group kept on 8 °C; (2) warm exposed 1 week and (3) warm-exposed 4 week. Warm-exposed groups were moved to 18 °C for 1 or 4 weeks, then moved back to 8 °C to induce sex. Throughout this experiment, polyps were fed and cleaned four times per week.

### Statistical analyzes

We used Generalized Linear Mixed-Models with a binomial distribution to analyze the effects of season on reproductive mode (sexual vs. asexual) and Linear-Mixed Models with a Gaussian distribution to analyze the effects of season on time to gonadogenesis after cooling in males and females. Time to gonadogenesis was log-transformed prior to analysis. We included strain ID as a random effect in all three models to take into account variation in the number of polyps per strain. Season, year, polyp age and type (wild-collected vs asexual descendant of a wild-collected polyp) were included as predictors. From the full model we excluded non-significant predictors in a stepwise manner based on largest p-values. Binomial GLMMs were implemented using the *lme4* package in R, while Gaussian LMMs were implemented using the *nlme* R package.

For the warm exposure experiments, we used Fisher’s exact test to analyze differences in the ratio of sexual and asexual individuals. No comparison on sex starting dates could be made because of the very low numbers of sexual individuals in some of groups (see Results).

## Results

### Establishment of field-collected strains

We established strains from N = 211 polyps (N = 54, 40, 59, 58, respectively for the four collection dates). The number of strains at the time of cooling was lower because of mortality: there were N = 182 (N = 50, 33, 53, 46) strains at the time of cooling. N = 194 (N = 40, 35, 53, 66) polyps died before lowering the temperature. N = 121 (N = 18, 31, 63, 9) polyps died after cooling or failed to produce any bud or gonad, therefore were excluded. Final sample size was N = 1243 polyps (N = 403, 204, 452, 184), with an average of 6.8 (8.1, 6.2, 8.5, 4.0) polyps per strain.

### Seasonal variation in sexual readiness

Season had a significant effect on reproductive mode and time to gonadogenesis in both males and females: strains established from autumn-collected polyps were significantly more likely to reproduce sexually in response to lowering the temperature (Table 1, Figure 1) and required less time for the production of gonads in both males and females (Table 1, Figure 2). In addition to season, polyp age was the only other factor influencing reproductive mode and time to gonadogenesis: older polyps were more likely to reproduce sexually and required less time to gonadogenesis (Table 1).

**Table 1.** Full and reduced models of the effects of season, polyp age, collection year and whether it was wild-collected or originated in the laboratory on reproductive mode (sexual vs. asexual), time to appearance of testes after cooling in males and time to appearance of eggs after cooling in females. All models contain strain ID as random factor.

| Predictor | Full model |  | Reduced model |  |
| --- | --- | --- | --- | --- |
|  | Estimate (SE) | t (P) | Estimate (SE) | t (P) |
| **Reproductive mode (binomial GLMM)** |  |  |  |  |
| Season (autumn vs. spring) | 2.231 ( 0.478 ) | 4.671 ( <0.001 ) | 2.237 ( 0.477 ) | 4.688 ( <0.001 ) |
| Age | 0.020 ( 0.006 ) | 3.204 ( 0.001 ) | 0.020 ( 0.005 ) | 3.801 ( <0.001 ) |
| Year (2019 vs. 2018) | -0.222 ( 0.417 ) | -0.531 ( 0.595 ) |  |  |
| Wild-collected (yes vs. no) | 0.048 ( 0.444 ) | 0.107 ( 0.915 ) |  |  |
| **Time to appearance of testes (Linear mixed model)** |  |  |  |  |
| Season (autumn vs. spring) | -0.414 ( 0.047 ) | -8.728 ( <0.001 ) | -0.401 ( 0.047 ) | -8.617 ( <0.001 ) |
| Age | -0.002 ( 0.001 ) | -4.063 ( <0.001 ) | -0.003 ( <0.001 ) | -6.461 ( <0.001 ) |
| Year (2019 vs. 2018) | -0.075 ( 0.046 ) | -1.613 ( 0.110 ) |  |  |
| Wild-collected (yes vs. no) | -0.049 ( 0.031 ) | -1.611 ( 0.108 ) |  |  |
| **Time to appearance of eggs (Linear mixed model)** |  |  |  |  |
| Season (autumn vs. spring) | -0.398 ( 0.053 ) | -7.535 ( <0.001 ) | -0.399 ( 0.046 ) | -8.643 ( <0.001 ) |
| Age | -0.003 ( 0.001 ) | -5.848 ( <0.001 ) | -0.003 ( <0.001 ) | -6.641 ( <0.001 ) |
| Year (2019 vs. 2018) | -0.004 ( 0.053 ) | -0.082 ( 0.935 ) |  |  |
| Wild-collected (yes vs. no) | 0.018 ( 0.031 ) | 0.580 ( 0.562 ) |  |  |

**Figure 1.**
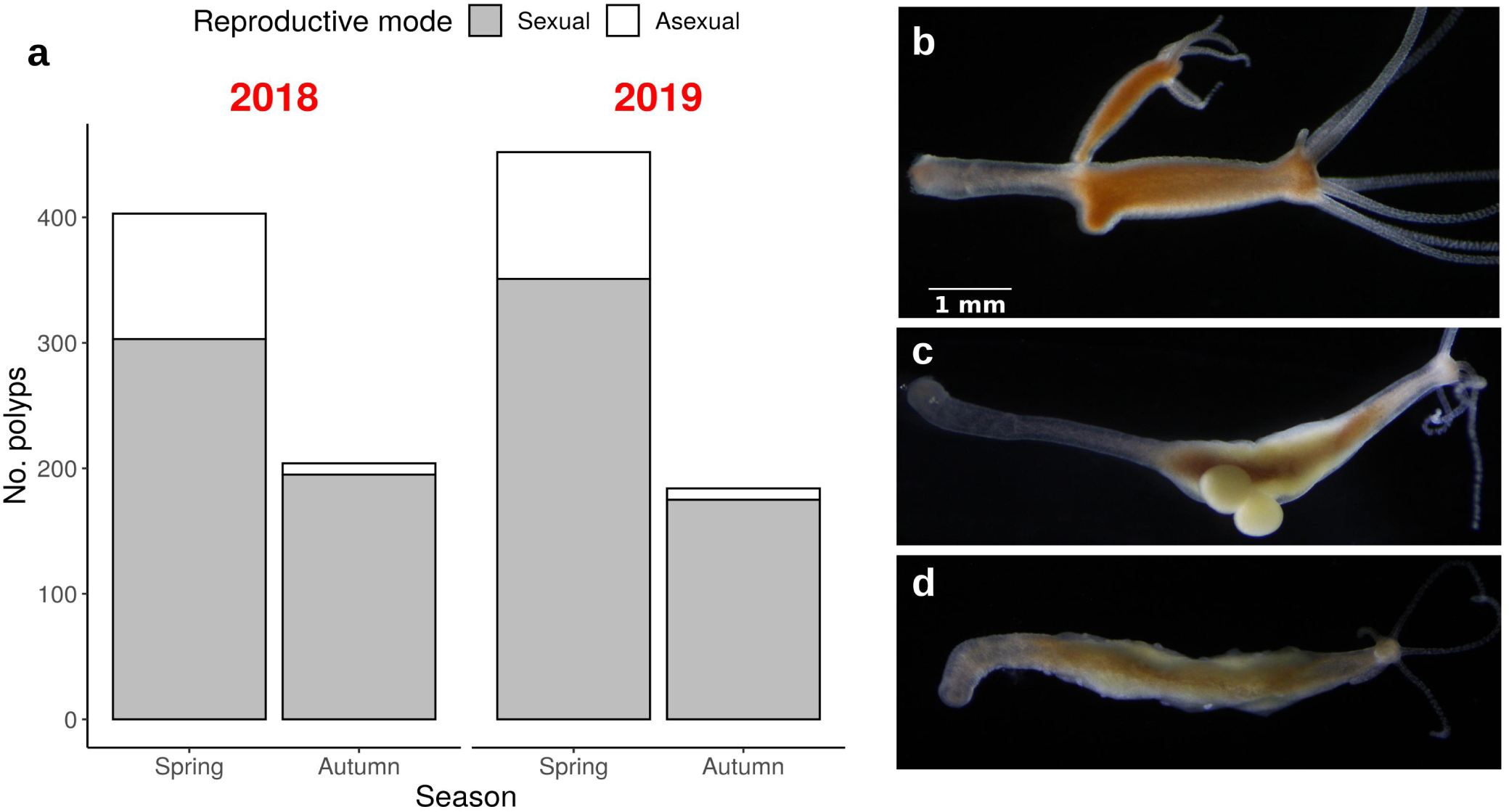
Proportion of sexual and asexual individuals in strains derived from spring- and autumn collected *H. oligactis* polyps (a) and photographs of asexual and sexual individuals (b: asexual, c: sexual female, d: sexual male)

**Figure 2.**
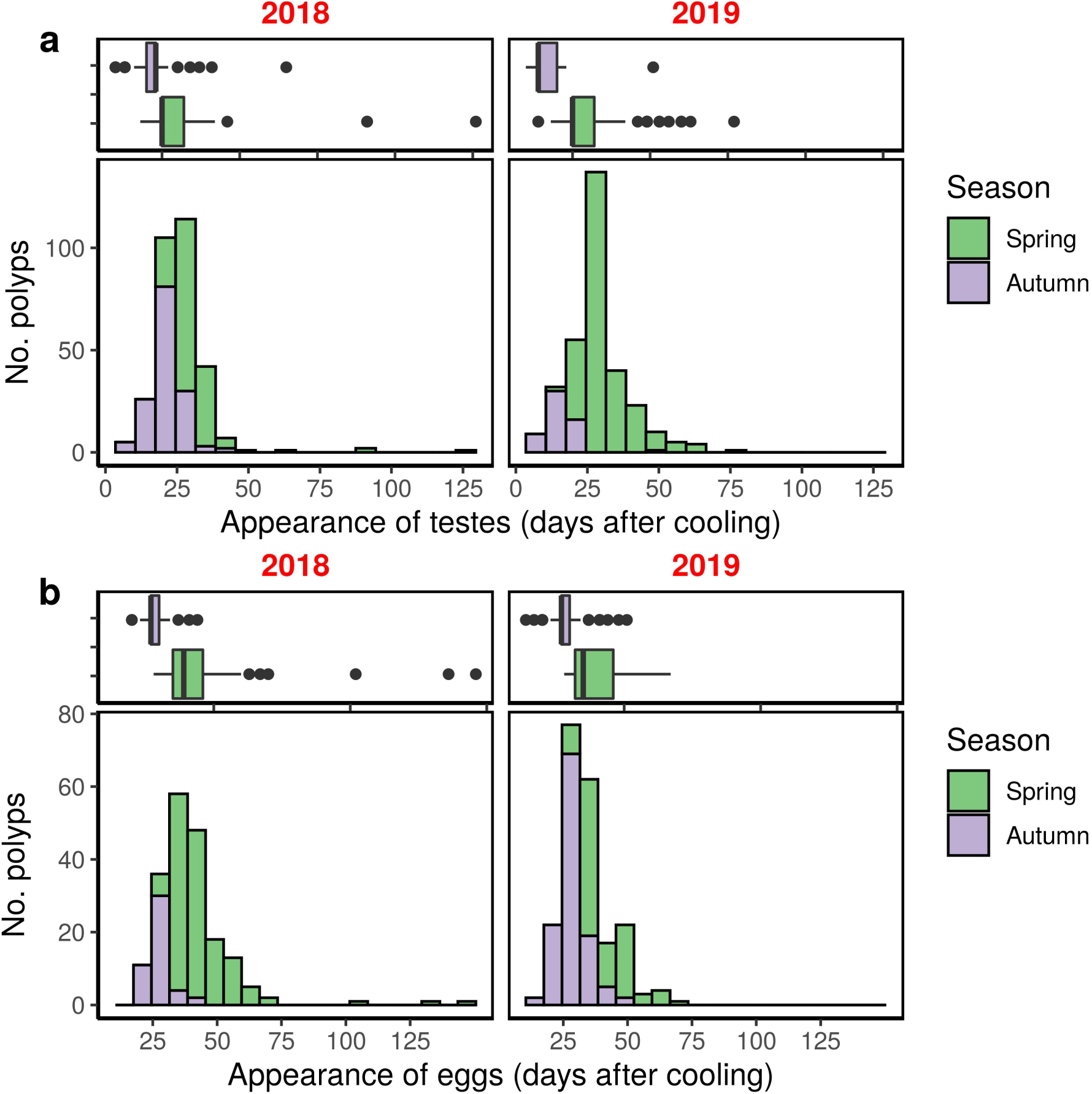
Time to initiate gonadogenesis in strains derived from spring- and autumn collected *H. oligactis* males (a) and females (b) kept in the lab under identical conditions. Data from two years (2018 and 2019) are shown separately

### Warm exposure experiment

We obtained N = 125 polyp in the female strain and N = 162 polyps in the male strain after 3 months of asexual propagation on 8 °C. N = 9 of these polyps died during the course of the experiment (N = 3 in the female strain, N = 6 in the male strain, all in the 4 weeks exposure group).

The proportion of sexual vs. asexual individuals was significantly different between the groups in both males and females (Fisher’s exact test p<0.001 for both sexes; Figure 3.). No sexual reproduction was observed in the group kept continuously on 8 °C. None of the polyps exposed to 1 week 18 °C underwent sexual reproduction in the male strain, but N=9 out of 41 polyps (22 %) in the female group initiated sex. The median starting day in this group was 63 days after cooling. In the groups exposed for 18 °C for 4 weeks, 31 out of 37 (84%) in the female strain and 40 out of 47 (85%) in the male strain underwent sexual reproduction. The median starting days were 31 and 40 days after lowering the temperature, respectively.

**Figure 3.**
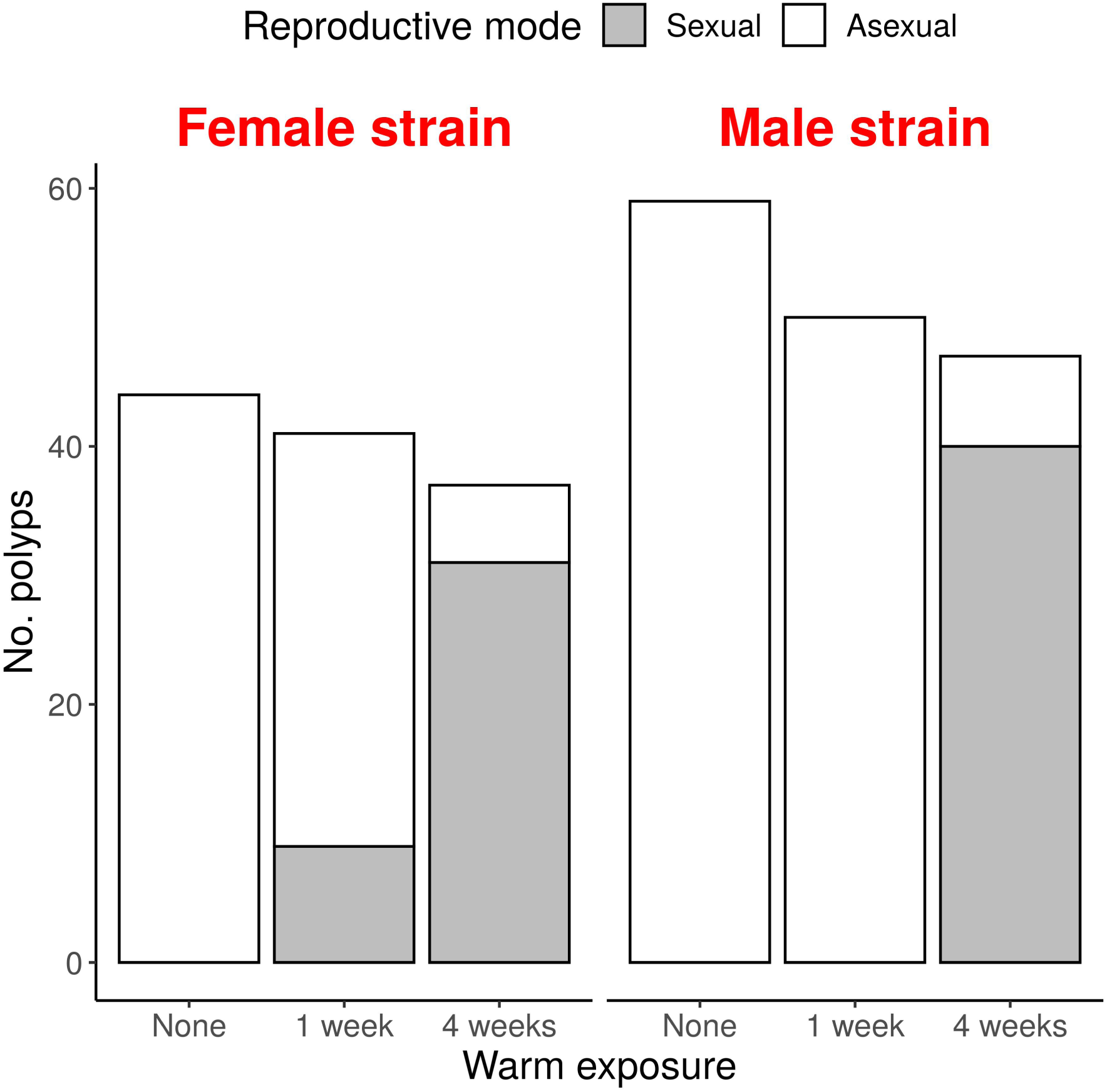
Effect of 1 or 4 weeks of warm exposure on the proportion of sexual individuals in a cold-acclimatized individuals of a male and female laboratory strains

## Discussion

Here we studied sexual readiness of *H. oligactis* strains established from spring- and autumn-collected polyps and kept under standard conditions in the laboratory. We found that the same environmental cue signaling the onset of winter (lowering the temperature from 18 to 8 °C) elicited different responses in the spring- and autumn collected strains. Sexual readiness was higher in strains derived from autumn-collected polyps: a higher proportion underwent sexual reproduction and it took significantly less time for them to start gonadogenesis.

In natural populations, sexual reproduction in *H. oligactis* occurs exclusively from late summer to early winter (Welch and Loomis 1924, Reisa 1973, Ribi et al. 1985, Sebestyén et al. 2018). Sexual reproduction results in the production of resting eggs that can survive the winter which the adults are thought to be less likely to tolerate due to freezing of water bodies and/or reduced food availability (Reisa 1973). Polyps that respond to temperature drop by switching to sexual reproduction might enjoy increased fitness because they can pass on their genes to the next generation before they die. A quick response to an environmental cue, on the other hand, might not be optimal if the drop in temperature is only transient and conditions favoring asexual reproduction return. An individual that invests into sexual reproduction under such conditions might receive a reduced fitness payoff compared to a strategy that delays sexual reproduction and maintains investing in clonal propagation throughout (Harvell and Grosberg 1988, Walsh 2013, Stelzer and Lehtonen 2016, Franch-Gras et al. 2017, Gerber et al. 2018).

Given the unpredictability of environmental conditions and the trade-offs associated with the decision to invest in diapausing forms, organisms are expected to be under selection to accurately time their life history decisions. One potential consequence of this selection is that overall sensitivity to environmental cues might change depending on predictability. For instance, if short-term deteriorations in the environment are frequent, sensitivity to environmental cues might decrease to avoid a switch to sexual reproduction when conditions favoring clonality might quickly return. Sensitivity to diapause-inducing environmental signals differs indeed greatly among closely related species and populations of facultatively sexual animals (e.g. Schröder and Gilbert 2004, Tessier and Cáceres 2004, Walsh 2013, Franch-Gras et al. 2017). Whether this variation is due to differences in the frequency of short-term deteriorations in the environment - and hence the reliability of environmental cues - is presently unclear.

An additional strategy for accurate timing in seasonal environments with unpredictability could be to increase sensitivity toward environmental cues with the progress of the seasons, such that investment into diapausing forms increases as the unfavorable season approaches. Examples of increased sensitivity to diapausing cues with seasons have been observed in water fleas and monogonont rotifers, which use transgenerational cues to infer the passage of time and ensure the optimal timing of resting egg production (Alekseev and Lampert 2001, Schröder and Gilbert 2004). Transgenerational maternal effects have been suggested as a general strategy to avoid diapause induction in insects during spring, when temperature and photoperiod are similar to autumn conditions (Reznik and Samartsev 2015, Tougeron et al. 2020). A similar pattern of increasing sensitivity with time was described for hydra strains kept in the laboratory: polyps had reduced propensity for sex after hatching from a resting egg, but sexual propensity increased during a period of three years in the laboratory (Noda 1982). Our observations complement those of Noda (1982) as we show that increased sexual readiness with the progress of seasons can be observed in wild-derived strains as well. This suggests that hydra polyps maintain low preparedness early in spring when the probability of the winter conditions is low,but increase preparedness when winter approaches.

In addition to describing the pattern of sexual readiness in spring- and autumn-collected hydra, our lab experiments also provided clues to the mechanisms underlying these differences. *H. oligactis* polyps generally respond to temperature drop by initiating sexual reproduction. However, there is variation in this response: younger polyps have reduced propensity for sex, lower fecundity and a higher post-reproductive survival rate (Sebestyén et al. 2019). Previously, we observed that polyps surviving after sexual reproduction are unlikely to undergo sexual reproduction again, if they are continuously maintained on 8 °C (Tökölyi J., pers. obs.). In this study, we now formally show that sex does not occur in animals propagated asexually on 8 °C (up to several months). This asexual propagation phase is likely to be part of the natural life cycle of the species at the end of the winter, as asexual polyps persist in the population after sexual reproduction and can reach high population density (Welch and Loomis 1924; J. Tökölyi, pers. obs.). Together, these observations suggest that cold exposure alone is not sufficient to induce sex in this system.

Since a drop in temperature, but not continuous cold exposure appears to be crucial to induce sex in *H. oligactis*, we performed a simple lab experiment to see how the reciprocity of cold and warm periods affects sexual readiness. In this experiment, we simulated seasonal changes in temperature: we first lowered temperature then increased it to 18°C for 1 or 4 weeks in the treated groups, while controls remained on cold. Finally, we returned warm-exposed hydra to 8 °C to attempt initiating sexual reproduction. The results of this experiment show that sexual reproduction is very low in the group exposed for 1 week to warm, but was much higher in the group that received a 4 week warm exposure. Overall, this experiment suggests that warm exposure increases sexual readiness and that reciprocal changes in cold and warm periods are required to elicit sexual reproduction in this species. Reliance on such reciprocal temperature changes might ensure the correct timing of sex, since it always will occur when a cold period follows a longer warm period (i.e. autumn, as normally observed in the natural habitats of this species). While the warm periods in our experiment were not as long as normally observed in the natural habitats of these strains, they nevertheless capture the pattern of seasonal fluctuations normally experienced by them.

From a proximate perspective, sex determination and the sexual phenotype in hydra is dependent on a population of germline stem cells (reviewed in Nishimiya-Fujisawa and Kobayashi 2018). These germline stem cells can be either male and female and derive from a common stock of multipotent interstitial stem cells that also give rise to somatic cell types (e.g. nematocytes, nerve cells; Nishimiya-Fujisawa and Kobayashi 2018). In *H. oligactis*, it is well established that the differentiation of germline stem cells to gametes occurs at temperatures below 12 °C (Littlefield 1991, Littlefield et al. 1991). Much less is known, however, about the factors that influence the proliferation of germline stem cells or their differentiation from multipotent interstitial stem cells. Our observations lead us to hypothesize that germline stem cells might be absent or in very low numbers in the cold asexual strains. First, polyps kept on 8 °C continuously did not undergo sex, but reproduced asexually continuously. Second, a short exposure to 18 °C was not sufficient to elicit sex, only a small percentage in the female strain initiated gonadogenesis. By contrast, the same drop in temperature after a longer warm exposure was sufficient to elicit sex with high probability. A plausible explanation for the pattern observed by us could be that the germline stem cells in this species require high temperatures to develop and/or proliferate and a 1 week exposure to 18 °C is not enough for the accumulation of germline stem cells. Testing this hypothesis will require quantification of germline stem cells under different temperature regimes.

To summarize, we have shown that in *H. oligactis*, a facultatively sexual freshwater cnidarian with diapausing eggs, sensitivity to an environmental cue that elicits sexual reproduction changes during the seasons. Polyps appear to maintain low sexual readiness during the spring, but sexual readiness increases as onset of unfavorable period approaches. These observations complement a previous study by Noda (1982) demonstrating similar effects under lab conditions, and studies in other facultatively sexual species (monogonont rotifers and *Daphnia*; Alekseev and Lampert 2001, Schröder and Gilbert 2004) documenting seasonal changes in sensitivity to diapause-inducing environmental cues. We complemented our observations of field-collected strains with experiments in lab strains and these suggest that the increase in sensitivity in hydra could be explained by long-term exposure of polyps to high temperature. The exact developmental mechanisms underlying this phenomenon remains to be explained in the future. The reliance on both cold and warm periods to complete the sexual / asexual cycle in *H. oligactis* could ensure proper timing of sex in this species. However, this reliance is likely to have a major impact on future adaptation of this species to global warming, the implications of which are presently hard to predict.

## Acknowledgements

This study was supported by NKFIH grant FK 124164. JT and MM were supported by the ÚNKP New National Excellence Program of the Ministry for Innovation and Technology (ÚNKP-19-3 and ÚNKP-19-4). JT was supported by a János Bolyai Research Scholarship of the Hungarian Academy of Sciences. We are grateful to the following people for lab assistance: Berta Almási, Beatrix Kozma, Dávid Tenkei, Erzsébet Ágnes Nehéz, Milla Makó and Flóra Sebestyén.

